# Analysis of physiological noise in quantitative cardiac magnetic resonance

**DOI:** 10.1101/581496

**Authors:** Terrence Jao, Krishna Nayak

**Affiliations:** Department of Biomedical Engineering, Viterbi School of Engineering, University of Southern California, Los Angeles, California, United States of America; Ming Hsieh Department of Electrical and Computer Engineering, Viterbi School of Engineering, University of Southern California, Los Angeles, California, United States of America

## Abstract

**Purpose:** To determine the impact of imaging parameters on the temporal signal-to-noise ratio (TSNR) of quantitative cardiac magnetic resonance (MR) in humans, and to determine applicability of the physiological noise covariance (PNC) model for physiological noise (PN).

**Methods:** We conducted MRI experiments in four healthy volunteers, and obtained series of short-axis cardiac images acquired with snapshot balanced steady-state free precession (bSSFP) and snapshot gradient echo (GRE) using a broad range of spatial resolutions and parallel imaging acceleration factors commonly used in quantitative cardiac MR. We measured regional SNR and TSNR in these datasets and fit the measurements to the PNC model for PN, which assumes that PN scales with signal strength.

**Results:** The relationship between SNR and TSNR in human cardiac MR without contrast preparation was well modeled by the PNC model. SNR consistently decreased as the spatial resolution (matrix size) and acceleration factor (R) increased for both GRE and bSSFP imaging. TSNR varied linearly with SNR using GRE imaging, when SNR was low (SNR < 20), and approached an asymptotic limit using bSSFP imaging, when SNR was high (SNR > 40).

**Conclusions:** The PNC model can be used to guide the choice of matrix size and acceleration factor to optimize TSNR in stable contrast cardiac MR, such as T2-prepared Blood-Oxygen-Level-Dependent (BOLD) and several variants of Arterial Spin Labeled (ASL) cardiac MR.

## Introduction

Quantitative cardiac magnetic resonance (MR) has shown clinical value in a wide array of applications, including T1, T2, and ECV mapping as well as first-pass and non-contrast perfusion imaging. Quantitative cardiac MR maps are formed from multiple images, which make them susceptible to errors caused by signal fluctuations of physiological origin, such as cardiac or respiratory motion. This is known as physiological noise (PN) and impacts the temporal signal-to-noise (TSNR) ratio which ultimately determines measurement variability.

Recent advances in cardiac MR have enabled clinically feasible quantitative mapping of magnetic relaxation properties and perfusion of the myocardium (1). T1 and ECV mapping can assess diffuse myocardial fibrosis and infiltration (2–4) and can measure extracellular volume when acquired with contrast (5,6). Inflammation and edema in the heart can be detected with T2 mapping (7) while T2* is sensitive to iron induced field inhomogeneity in hemochromatosis (8) and hemorrhage (9). Quantitative myocardial perfusion imaging performed using either first-pass cardiac MR (10), arterial spin labeling (ASL) (11), or blood oxygen level dependent (BOLD) (12) has been shown to detect regions of ischemia to evaluate coronary artery disease severity. These quantitative maps are all formed by imaging the myocardium multiple times with different magnetization preparations to vary the sensitivity of each image to the parameter of interest. These preparations produce changes in the signal intensity of the acquired images, which are fit to models of the underlying physiology or relaxation parameter to generate a quantitative map.

Unexpected signal changes that are not from the designed preparations will cause errors in the parameter estimates. These errors arise from thermal noise as well as physiological fluctuations from cardiac, respiratory, and hemodynamic motion. Thermal noise is well understood and is modeled as an independent additive bivariate Gaussian process. It can be easily measured from noise-only data (acquired without RF excitation) or from a manually-selected noise-only region within an image. In contrast, the physiological noise distribution is largely unknown and cannot be directly measured. Physiological noise contributes to the temporal standard deviation of an image series, and can become a dominant contributor to measurement variability. In quantitative imaging, temporal signal-to-noise ratio (TSNR) is measured as the mean signal intensity within a region-of-interest (ROI) divided by its temporal standard deviation.

Triantafyllou et. al. (13) demonstrated that in the brain image acquisition parameters, including spatial resolution, parallel imaging acceleration factor, and flip angle, altered the TSNR. They were also able to experimentally confirm that physiological noise increases proportionally with the signal strength (14). This is problematic because a simple increase of signal strength through higher field or denser receiver arrays may not translate to improvements in TSNR if physiological noise is dominant. In this case, efforts should be made to increase spatial resolution or speed acquisition through parallel imaging. Northrup et. al. (15) observed this phenomenon in ASL cardiac MR, which exhibited similar TSNR at 1.5T and 3.0T, despite the SNR advantage at 3.0T. Do et. al. (16) used parallel imaging to speed up the acquisition of ASL cardiac MR and observed an increase in TSNR despite a decrease in SNR.

In this study, we measure the SNR and TSNR of cardiac mid short-axis image series under different spatial resolutions and parallel imaging acceleration factors and assess the ratio between physiological and thermal noise. We study snapshot bSSFP and GRE acquisitions because they are the most commonly used acquisition schemes for cardiac MR T_1_, T_2_, and ECV mapping and for myocardial perfusion imaging. While the results of this work are specific to our MRI apparatus, receiver coil, and pulse sequence, it is simple to reproduce and need only be performed once. To the best of our knowledge, this is the first study that experimentally examines how the choice of imaging parameters affects TSNR and *σ*_*P*_ for quantitative cardiac MR.

## Materials and Methods

### Noise Analysis

Our analysis of physiological noise and TSNR follows the work of Triantafyllou et. al. (13) for multi-channel coil arrays in fMRI image series in the brain. The total noise, *σ* in a cardiac MR image series is modeled as the independent sum of thermal noise, *σ*_*t*_ and physiological noise, *σ*_*p*_.

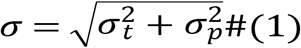

Physiological noise is described by the Krüger and Glover model (PNC) (14) to scale proportionally by a constant, *λ*, with the MR signal level, *S*, such that *σ*_*p*_ = *λS*. According to this model for noise, TSNR can be derived from image SNR as follows:

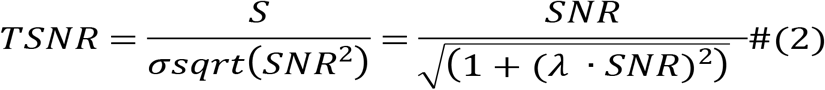

Recently, Triantafyllou et. al. proposed a new model that captures coil to coil noise correlations from physiological fluctuations by splitting the noise covariance matrix into a thermal and physiological component (17). This model relaxes the assumption that PN must scale with the signal level and reduces to the KG model when the physiological noise covariance matrix scales with the signal level. The correlations of physiological noise sources that are not signal level dependent are described by an additional parameter, *α*, to generalize the model such that

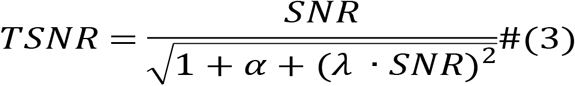

In practice, both *λ* and *α* are unknown and TSNR is directly measured by dividing the mean signal intensity by its temporal standard deviation. However, an important result of this model is that TSNR reaches an asymptotic limit of 1/*λ* when SNR becomes large. This implies that attempts to increase TSNR by increasing SNR will fail if SNR is already high.

From measurements of SNR and TSNR, the ratio of physiological noise to thermal noise can be measured as follows:

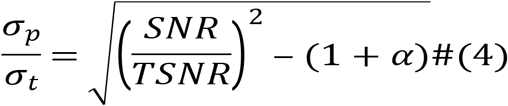

This noise ratio determines whether an image series is thermal noise or physiological noise dominant. When images are thermal noise dominant, improvements in TSNR can be achieved by increasing the signal, through higher field strength of denser surface coils. However, when images are physiological noise dominant, improvements in SNR have negligible impact on TSNR. Instead, SNR can be traded for other gains, such as in spatial resolution or parallel imaging acceleration factor up until the physiological noise is comparable to thermal noise.

### Experimental Methods

Images were acquired on a 3T GE Signa Excite HD with an 8-channel cardiac receiver array in 4 healthy volunteers. Written informed consent was acquired from all volunteers using a protocol approved by the institutional review board. Images were acquired using snapshot bSSFP and GRE. All images were acquired during mid-diastole at a single mid short-axis slice. For each unique set of acquisition parameters, twenty images were acquired. Images were acquired with a 1 second breath hold with a 3 second wait between acquisitions to allow the magnetization to fully recover. A cardiac cinema/video [CINE] scout scan was acquired in each subject to determine the exact timing of mid-diastole relative to the ECG R-wave. Noise only images without RF were also acquired to calculate the thermal noise and coil noise covariance matrix in each subject.

In each subject, data were acquired at various matrix sizes (192 × 96, 192 × 128, 192 × 192) and accelerated (1, 1.33, 1.6, 2) by GRAPPA (18) and partial Fourier imaging. Other imaging parameters were fixed throughout the experiment, which include TR/TE of 3.6/1.8 ms, FA of 50^0^/5^0^ for <u>b</u>SSFP and GRE respectively, BW of ±125 kHz, and slice thickness of 10 mm. This led to a total of 24 different imaging settings (3 matrix sizes, 4 acceleration factors, 2 imaging sequences). Differences in the size of the volunteers led to different choices of FOV and spatial resolution. Specific imaging parameters are summarized in Table 1, and inter-subject variations in the FOV and spatial resolutions are summarized in Table 2.

**Table 1:** Acquisition parameters used for acquiring cardiac image series at various resolutions and acceleration factors. *Imaging settings at 192×192 and 192×144 have long imaging windows that occupy up to half of the cardiac cycle.

| Reconstructed Matrix | Acquired Matrix | Partial Fourier | Acceleration | Imaging Window (ms) |
| --- | --- | --- | --- | --- |
| **192 x 96** | $192 \times 96$ | 1 | 1 | 336 |
| | $192 \times 72$ | 6/8 | 1.33 | 252 |
| | $192 \times 60$ | 1 | 1.6 | 210 |
| | $192 \times 48$ | 6/8 | 2 | 168 |
| **192 x 128** | $192 \times 128$ | 1 | 1 | 448 |
| | $192 \times 96$ | 6/8 | 1.33 | 336 |
| | $192 \times 76$ | 1 | 1.6 | 266 |
| | $192 \times 60$ | 6/8 | 2 | 210 |
| **192 x 192** | $192 \times 192$ | 1 | 1 | 672* |
| | $192 \times 144$ | 6/8 | 1.33 | 504* |
| | $192 \times 108$ | 1 | 1.6 | 378 |
|  | 192 × 84 | 6/8 | 2 | 294 |

**Table 2:** Variations in FOV and spatial resolution in the different subjects. Resolution is reported for matrix sizes of 192 × (96, 128, 192).

| Subject | FOV (mm<sup>2</sup>) | Resolution (mm) |
| --- | --- | --- |
| 1 | 320 | 1.67 × (3.33, 2.50, 1.67) |
| 2 | 360 | 1.88 × (3.75, 2.81, 1.88) |
| 3 | 360 | 1.88 × (3.75, 2.81, 1.88) |
| 4 | 320 | 1.67 × (3.33, 2.50, 1.67) |

### Image Reconstruction and Data Analysis

Image were reconstructed offline using software written in Matlab (Mathworks Inc. Natick, MA, USA). Partial Fourier image reconstruction was carried out using homodyne reconstruction with a linear ramp weighting while GRAPPA reconstruction was performed using a custom implementation. Images were coil combined using an optimal B1 weighted combination (19) and aligned with non-rigid motion correction using advanced normalization tools (ANTs) (20).

SNR maps were calculated for each image using the “pseudo-replica” method described by Robson et al. (21) For each image, correctly scaled and correlated synthetic noise is added to acquired k-space data before reconstruction. This is repeated multiple times, each with a different instantiation of synthetic noise, to generate an image stack. Under the assumption that the reconstruction process acts independently on the signal and the noise, SNR can be calculated as the ratio of the image reconstructed without noise addition to the standard deviation of the image stack. In contrast, TSNR was measured from the “actual multiple replicas” that were acquired. It is calculated as the mean intensity of a pixel or ROI of the left ventricular myocardium divided by the temporal standard deviation. The SNR and TSNR for each of the 24 individual imaging parameters were averaged across the four subjects, and fit to the PNC model of Eq 3 using a non-linear least squares algorithm provided by Matlab to estimate the physiological noise scaling parameter, *λ* and physiological coil covariance parameter, *α*. The noise ratio from Eq 4 was subsequently calculated from SNR, TSNR and *α*.

## Results

Figure 1 shows a representative SNR, TSNR, and noise ratio map from one volunteer with an acceleration factor of 1 using a 192 × 192 matrix. SNR is always greater than or equal to TSNR. The noise ratio map is low in areas with little movement such as the musculature in the back and highest in areas with large fluctuations in signal intensity, such as blood vessels.

**Figure 1:**
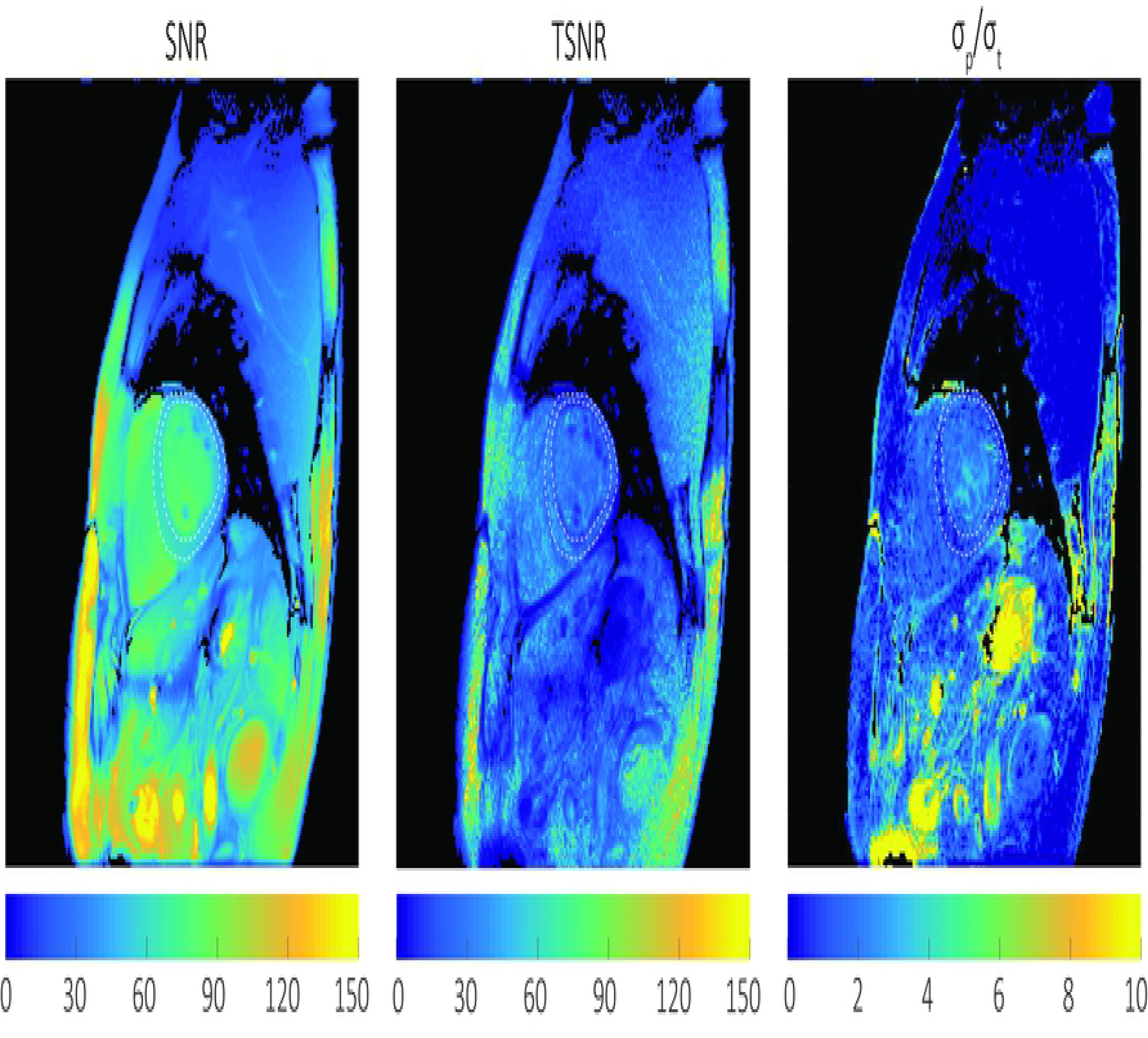
Representative SNR, TSNR, and *σ*_*P*_/ *σ*_*T*_ maps from a healthy volunteer. Images were acquired with a matrix size of 192×192, R= 1, Resolution = 1.67×1.67mm^2^. The SNR, TSNR, and *σ*_*P*_/*σ*_*T*_ ratio within the left ventricular myocardium ROI (white dashed line) was 51.3± 9.8, 30.5±10.2, and 1.49 ± 0.56 respectively. The noise ratio map is lowest in areas with little movement, such as the muscles of the back and highest in areas with large fluctuations in signal intensity such as blood vessels. In a static phantom and stable imaging environment, TSNR = SNR. In vivo, TSNR ≤ SNR due to additional sources of *σ*_*P*_. The *σ*_*P*_/*σ*_*T*_ ratio determines whether *σ*_*P*_ or *σ*_*T*_ is the dominant source of noise. When *σ*_*P*_ is dominant (*σ*_*P*_/*σ*_*T*_ >> 1), SNR gains will not improve TSNR.

Figure 2A shows TSNR as a function of SNR in the left ventricular myocardium. Each point represents an average across subjects while the error bars represent the standard deviation within the ROI averaged across subjects. Red solid points represent bSSFP acquisitions while blue outlined points represent GRE acquisitions. The points were fit to Eq 3, shown as the solid black line, with a physiological noise scale factor, *λ*, of 0.0197. The lower and upper 95% confidence interval of *λ* was 0.0173 and 0.0221 and are plotted with their corresponding dotted black lines. The physiological noise covariance factor, *α*, is 0.817 with 95% confidence interval of 0.383, 1.252. The vertical dashed red line corresponds to the SNR in which the noise ratio is unity and has a value of 50.8 (1/*λ*). The noise ratio is less than unity when left of the line, and greater than unity when right of the line. The horizontal dashed red line corresponds to the asymptotic value for TSNR which also has a value of 50.8 (1/*λ*).

**Figure 2:**
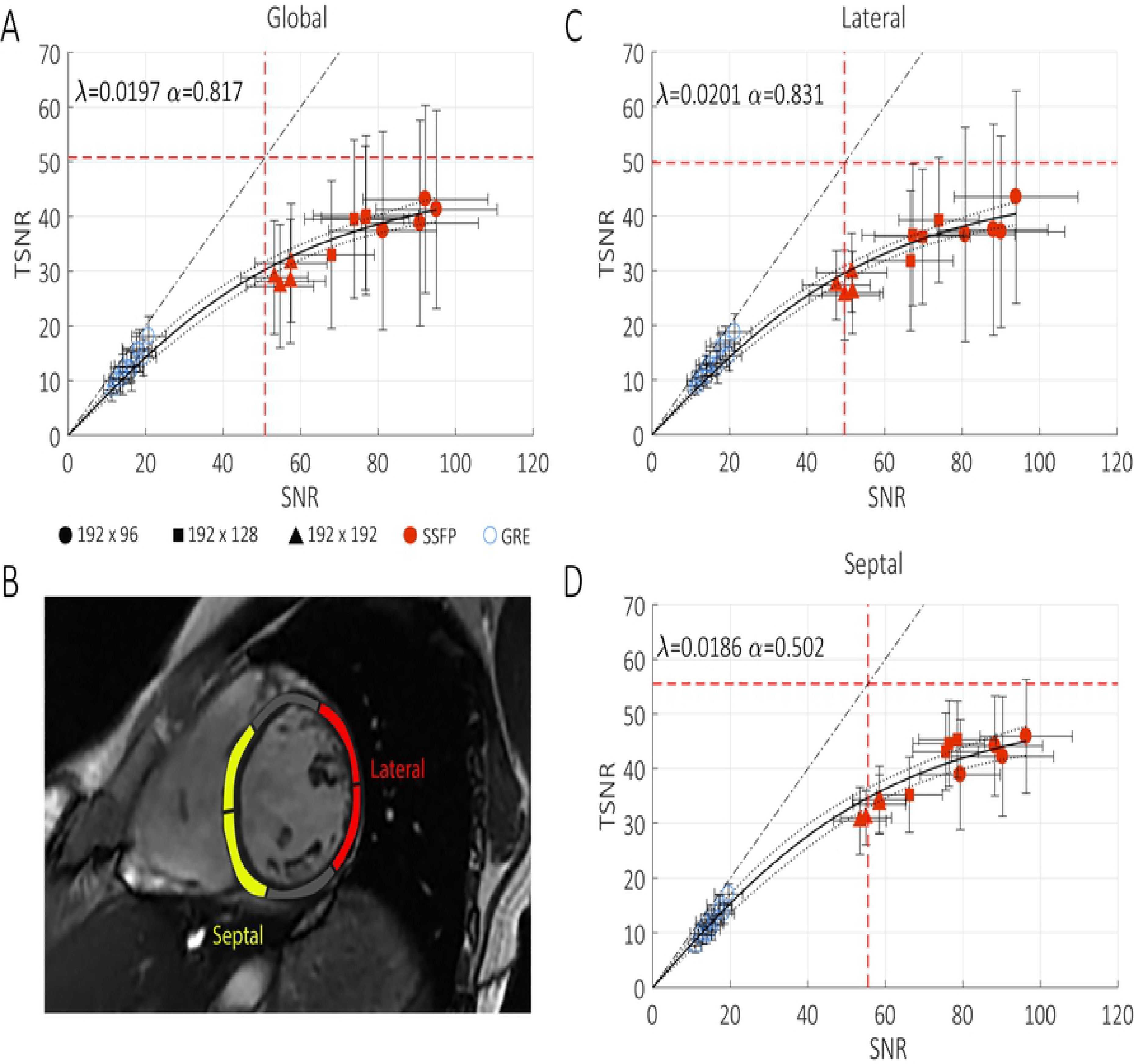
Plots of temporal signal-to-noise ratio (TSNR) versus signal-to-noise ratio (SNR). Separate plots are for A) all LV myocardium, C) septal myocardium and D) lateral myocardium, from mid-short axis slices. Septal and lateral segments are illustrated in B). Each point represents an average over four subjects while the error bars represent the standard deviation within the ROI averaged over four subjects. bSSFP acquisitions are shown in solid red while GRE acquisitions are shown in outlined blue. The solid black line represents the fit to the PNC model along with the lower and upper 95% confidence interval shown in dotted black. The dashed black line indicates the line of unity. Vertical and horizontal dashed red lines represent when the noise ratio is equal to one and the asymptotic limit of TSNR respectively. Both occur at 1/*λ*. This indicates that the measured TSNR versus SNR tradeoff is well characterized the PNC model. At low SNR, TSNR varies linearly with SNR while at high SNR, TSNR approaches an asymptotic limit. Note that variations of SNR and TSNR within the lateral myocardium were 1.58 and 1.41 times greater than variations within the septal myocardium. The lateral myocardium also exhibited higher physiological noise (*λ* = 0.0201, *α* = 0.831) and hence lower asymptotic TSNR than the septum (*λ* = 0.0186, *α* = 0.502).

The data were also analyzed within the lateral and septal walls, which are shown in Figure 2C and Figure 2D respectively. Septal and lateral segments are defined by the AHA 17-segment model (22) and are shown in Figure 2B. SNR and TSNR variation within the lateral wall is higher than in the septum using bSSFP imaging as indicated by the error bars. The standard deviation for SNR and TSNR measurements are 12.88 and 11.75 respectively within the lateral wall and are 7.67 and 9.26 respectively within the septum. *λ* is higher within the lateral walls with a value of 0.0201 with lower and upper 95% confidence interval of 0.0179 and 0.0244 while *λ* is lower in the septum with a value of 0.0186 with confidence interval of 0.0162 and 0.0209. This leads to an asymptotic value for TSNR of 49.7 and 53.9 for the lateral wall and septum respectively. *α* within the lateral and septal walls are 0.831 with 95% confidence of 0.446 to 1.216 and 0.502 with 95% confidence of 0.103 to 0.900 respectively.

Figure 3 contains plots of average SNR, TSNR, and *σ*_*p*_/*σ*_*t*_ within the left ventricular myocardium ROI as a function of the image matrix size and acceleration factor. SNR consistently decreased as the spatial resolution (matrix size) and acceleration factor (R) increased for both GRE and bSSFP imaging. In GRE imaging, TSNR decreased as the spatial resolution and acceleration factor increased. The noise ratio was consistently lower than unity, represented by the dashed red line, for all matrix sizes and acceleration factors. In contrast, TSNR only slightly decreased as the matrix size increased in bSSFP imaging, but remained constant when acceleration factor increased. The noise ratio in bSSFP imaging was consistently above unity and only decreased to approach unity at the largest matrix size and acceleration factor.

**Figure 3:**
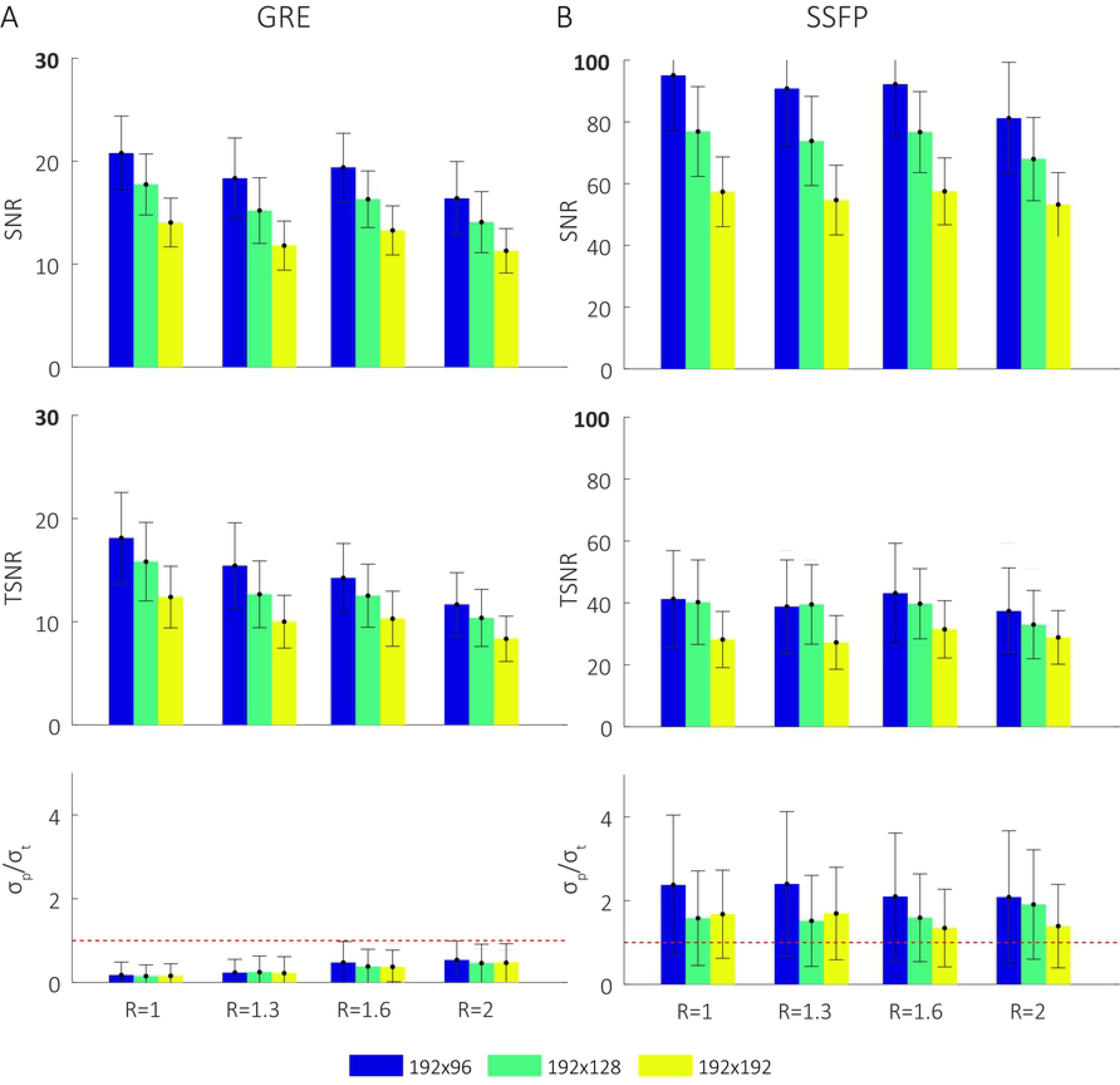
Comparison of SNR, TSNR, and the *σ*_*p*_/*σ*_*t*_ for several image matrix sizes and acceleration factors. A) GRE and B) bSSFP. As expected, SNR decreases with increasing matrix size and higher acceleration factors. In GRE, TSNR follows a similar trend and the noise ratio is < 1 for all imaging settings. In bSSFP, TSNR remains relatively constant and the noise ratio is ≥1 at low acceleration factors and approaches 1 only at higher accelerations and larger matrix sizes.

## Discussion

In this work, we investigated how changes in spatial resolution and parallel imaging acceleration factor affect SNR, TSNR, and the noise ratio for quantitative cardiac MR using snapshot GRE and bSSFP imaging. We found that when spatial resolution is made coarser and acceleration factor is reduced, SNR consistently increases while TSNR faces diminishing returns. In particular, we observed that TSNR varies linearly with SNR using GRE imaging (SNR < 20) and approaches an asymptotic limit using <u>b</u>SSFP imaging (SNR > 40). This makes the noise ratio (*σ*_*p*_/*σ*_*t*_) thermal noise dominant (<1) for GRE and physiological noise dominant (>1) for bSSFP in all but the largest of matrix sizes and acceleration factors (192 × 192, R=2). These results are predicted by the PNC model; physiological noise scales with signal strength and above a certain threshold, further improvements in SNR do not necessarily translate to improvements in TSNR.

The relationship between SNR and TSNR, as descried by the PNC model, can be categorized into three regimes - the linear, transition, and asymptotic regime. The linear regime is defined as the range of SNR where the noise ratio is less than unity while the asymptotic regime is defined as the range of SNR where TSNR is greater than 95% of its asymptotic value. The transition regime lies between the two. One should avoid imaging in the linear regime because TSNR can always be improved by increasing SNR. Similarly, one should avoid imaging in the asymptotic regime because SNR can be sacrificed with little to no loss in TSNR, which will yield improvements in spatial resolution or scan time. Quantitative cardiac MR protocols should be performed in the transition regime, and the experiments described in this manuscript can inform appropriate parameter selection.

In the original KG model, physiological noise can be interpreted as a modulation of signal intensity within a voxel caused by complex physiological processes, such as pulsatile blood flow or the effect of deoxyhemoglobin in the BOLD effect. These effects are local and occur within the voxel. In contrast, with bulk motion, tissue signal intensities are modulated as they move through the spatial gradient of the receiver sensitivity profile over time and can no longer be accounted for by the KG model. In cardiac imaging, the major sources of PN are bulk motion during the cardiac cycle and respiration. The PNC model relaxes the assumption that PN must scale with the signal level and can be used to model the major sources of PN in CMR. While the PNC model was originally developed for long-TR EPI imaging used in BOLD fMRI, we found that it is applicable to snapshot GRE and bSSFP often used for cardiac imaging. Snapshot imaging is performed in a transient state and is designed to capture the longitudinal magnetization prior to imaging.

Previous studies have only investigated the relationship between SNR and TSNR for functional MRI of the brain (13,14). In those studies, reported values for the physiological noise scaling parameter, *λ*, (0.009) are significantly lower than values we report for the heart (0.0197). This is likely due to cardiac and respiratory motion found in the heart that are absent in the brain. We also reported regional variations in SNR and TSNR and found that *λ* is 1.12 times larger in the lateral wall (0.0201) than in the septum (0.0186). Similarly, alpha is 1.66 times larger in the lateral wall (0.831) than in the septum (0.502). We suspect that this is due to greater off-resonance sensitivity and motion found in the lateral wall.

One concern about the results of this study is the precision of the TSNR estimate, which is dependent on the accuracy of the noise standard deviation. The noise standard deviation is determined by the number of images acquired and pixels within the ROI. Using a Monte Carlo simulation for 20 images with 150 pixels within the left ventricular myocardium, 50 pixels for regional analysis, and individual pixel-wise analysis, the relative error of the TSNR measurement was 1.6%, 2.1%, and 12.97% respectively. This is below the inter-subject variation for SNR and TSNR and does not alter the conclusions of the study. Another concern in our analysis is the existence of physiological noise within an imaging window, which may be more pronounced in longer acquisition windows (e.g. 672 ms for a 192 × 192 matrix size without acceleration). TSNR can only be measured between images and all noise within an image is assumed to come from thermal noise. Therefore, SNR may be underestimated and the SNR-TSNR curve may be shifted to the right. Rapid acquisition schemes such as EPI or spiral imaging may be evaluated to observe the degree of this bias. We hypothesize that these acquisition schemes, with their shorter imaging window, may even lie on a completely different SNR-TSNR curve with a lower *λ* and higher asymptotic TSNR.

A natural follow-up to this study would be to investigate whether the relationship between SNR, TSNR, and the noise ratio changes in systolic imaging, during exercise or pharmacologic stress, or with different cardiac pathologies. For instance, arrhythmia and stress may cause more cardiac motion during imaging and decrease TSNR while heart failure and post-acute myocardial infarct scar may reduce cardiac motion and increase TSNR. These studies may be difficult to perform in humans, but could potentially be examined in a large animal model, where respiration can be controlled and many images can be acquired. The drawback is that animals are typically anesthetized and may have a different stress response than humans. Nevertheless, it would be insightful to determine whether image acquisitions parameters can be optimized for cardiac phase, stress, or pathology.

Another limitation of this study is that we only considered equilibrium contrast. Quantitative cardiac MR can involve large variations in contrast due to magnetization preparation and/or passage of contrast agents. A single experiment (i.e. T1 mapping or T2 mapping) can have images with a broad range of SNR and regional contrast. While changes in image acquisition parameters are expected to cause shifts along the SNR vs. TSNR curve, changes in contrast or contrast preparation parameters could cause shifts of the entire curve. The findings of this study are therefore only directly applicable to quantitative cardiac MR techniques that utilize snapshot imaging and have stable contrast between images. This includes T2-prepared BOLD cardiac MR (23) and ASL cardiac MR using PICORE, EPISTAR, or steady-pulsed labeling (24).

In summary, we demonstrated that the SNR and TSNR of equilibrium snapshot cardiac MR images can be parametrized by the PNC model and varies with different image acquisition parameters, including spatial resolution and acceleration factor. However, we have also identified additional imaging parameters that must be considered for cardiac imaging, the precise timing and duration of imaging within the RR interval. Future work needs to be done to characterize these different regimes for systolic imaging, stress imaging, and in different cardiac pathologies.

## Acknowledgements

We thank our team of researchers at the Magnetic Resonance Engineering Laboratory including Hung Phi Do, Vanessa Landes, Ahsan Javed, Yongwan Lim, Yannick Bliesener, and Xin Miao.

